# Antibacterial activity of total flavonoids from *Ilex rotunda Thunb*. and different antibacterials on different multidrug-resistant bacteria alone or in combination

**DOI:** 10.1101/457911

**Authors:** Yongji Wu, Beibei Chai, Lizhen Wang, Weijia Jiang, Mei Hu, Yuchuan Zhao, Hongbin Si

**Affiliations:** Institute of Animal Science and Technology of Guangxi University, Nanning, Guangxi 530004, China

**Keywords:** Antibacterial activity, Total flavonoids, *Ilex rotunda Thunb*, Multidrug-resistant bacteria, Synergistic, Additive

## Abstract

The problem of bacterial resistance is becoming more and more serious, which has become an urgent problem to be solved in human and veterinary. One approach to control and delay bacterial resistance is combination therapy in which antibiotics are given together with other antimicrobial or non-antimicrobial agents. Studies have shown that flavonoids from Traditional Chinese medicine (TCM) possess a high level of antibacterial activity against antibiotic resistant strains. The aim of this study was to evaluate the antibacterial effects of a combined therapy of total flavonoids from *Ilex rotunda Thunb*. and antibiotics against seven kinds of veterinary bacteria which were multidrug resistance bacteria. A microdilution checkerboard method was used to determine the minimal inhibitory concentrations of both types of antimicrobials, alone and in combination. The fractional inhibitory concentration index was calculated and used to classify observed collective antibacterial activity as synergistic, additive, indifferent or antagonistic.

From the performed tests, the total flavonoids and antimicrobial agents were combined to inhibit different multidrug-resistant bacteria, such as *Escherichia coli*, *Streptococcus*, *Pseudomonas aeruginosa*, *Enterococcus faecalis*, *Proteus vulgaris*, *Staphylococcus aureus*, *Acinetobacter baumannii*. For these bacteria, total flavonoids from *Ilex Rotunda Thunb*. presented synergistic or additive with different antibiotics and had a certain antibacterial effect on the separated multidrug-resistant bacteria. The study shows total flavonoids from *Ilex rotunda Thunb*. have potential as adjuvants for the treatment of animal bacterial diseases.

## 1. Introduction

With extensive use and abuse of antibiotics in both human and veterinary medicine, the problem of rapid spread of bacterial resistance is becoming more and more serious(Jones, Draghi, Thornsberry, Karlowsky, Sahm & Wenzel, 2004). So it is more difficult to find an effective drug treatment of animal bacterial diseases, the increase in dosage, development of drug resistance, drug residues and accumulation in body surely will do harm to body health, but also damage the ecological environment(Liu et al., 2017; Miravitlles & Anzueto, 2017).

China is fortune as it owns rich resources of traditional Chinese Medicine. Novel antibacterial action of plant extracts or effective bacteriostatic component, such as flavonoids, berberine, polysaccharides, saponin, volatile oil, etc. have been documented(Ramezani, Rahmani & Dehestani, 2017;Naz et al., 2017). Few plants extraction and effective bacteriostatic component exhibited synergistic or additive interaction with antibiotics against drug-resistant bacteria(Eskandary, Tahmourespour, Hoodaji & Abdollahi, 2017; Ye et al., 2017). Screening of crude extracts for synergistic or additive interaction with antibiotics is expected to provide bioactive compounds to be used in combinational therapy. These compounds may not have strong antibacterial activity but may enhance the activity of antibiotics synergistically or additively.

*Ilex rotunda Thunb*. belongs to the *Ilex* genus, the dried bark or root bark. The chemical constituents have been isolated from more than 20 kinds, mainly including three terpenoids, flavonoids, phenols, tannins and so on(Liu, Peng, Chen, Liu, Liang & Sun, 2017). The research on antibacterial, anti-inflammation is becoming more and more popular and accepted. Flavonoids are well-known antimicrobial compound against some pathogenic microorganisms and have been regard as one of potential sources of novel antimicrobial agents(Zakaryan, Arabyan, Oo & Zandi, 2017; Kurek, Nadkowska, Pliszka & Wolska, 2012). Flavonoids used alone or with some antibacterial drugs combination have certain antibacterial effects, and some antibiotics combined with antibacterial drugs can reduce the minimum inhibitory concentration of bacteria and antibiotic resistant strains to increase antibacterial activity(Barbieri et al., 2017; Usman et al., 2016).

In the present study, we evaluated synergistic and additivity effects of antibiotics administered at doses lower than their minimum inhibitory concentrations (MICs) with total flavonoids from *Ilex rotunda Thunb*. to test whether it would enhance the antibacterial activity of antibiotics to veterinary multidrug-resistant bacteria. Potential synergistic and additive antibacterial effects were quantified by the fractional inhibitory concentration index (FICI), which was determined using MICs obtained by the microdilution checkerboard method (Roy-Leon, Lauzon, Toye, Singhal & Cameron, 2005).

## 2. Materials and methods

### 2.1 *Preparation of total flavonoids from Ilex rotunda Thunb*

*Ilex rotunda Thunb*. were provided by Guangxi Taihua Pharmaceutical Corporation. The bark was dried in oven 40^°^C till constant weight achieved, then crushed into powder with a size smaller than No.30 mesh. Total flavonoids were extracted under the extraction conditions that ethanol concentration, 63.71%; solid-liquid ratio, 43.03ml/g; ultrasonic temperature, 54.01^°^C and ultrasonic power, 63.25% (443W). The extracts were collected, freeze-dried, ground and then stored at -20^°^C. For experiment purposes flavonoids extract was reconstituted in 5% dimethyl sulfoxide (DMSO) to a final concentration of 320mg/ml.

### 2.2 *Antimicrobial agents*

Amikacin, colistin, meropenem, sulfamonomethoxine, amoxicillin, mequindox, ceftriaxone sodium, cefotaxime sodium, ceftiofur sodium, ceftazidime, lincomycin, florfenicol, fosfomycin, rifampicin were obtained from Sigma-Aldrich (Shanghai). Amikacin, colistin, meropenem were prepared in sterile distilled water. Amoxicillin was dissolved in phosphate e buffer (PH=6.0, 0.01mol/L). Azithromycin was prepared in 95% ethanol and dissolved in broth. Ceftazidime was dissolved in sodium carbonate solution (NaCO3, the amount of anhydrous sodium carbonate is ten percent of ceftazidime). Rifampicin was dissolved in dimethyl sulfoxide (DMSO, final concentrations ranging from 0.0002% to 0.0003%). Sulfamonomethoxine was first dissolved in half of total volume hot water added a small amount of sodium hydroxide and then diluted with sterile distilled water to the working concentration.

### 2.3 *Bacterial strains and growth conditions*

*Escherichia coli*, *Streptococcus*, *Pseudomonas aeruginosa*, *Enterococcus faecalis*, *Proteus vulgaris*, *Staphylococcus aureus*, *Acinetobacter baumannii* were all obtained from the Chinese Veterinary Laboratory at Guangxi University (Nanning, China). The cultures were preserved at -20^°^C with 50% (v/v) glycerin solution at a ratio of 1:1 before use. Working cultures of 7 kinds bacterial were prepared from frozen stocks by 3 sequential transfers in 10ml liquid media (CAMHB for *Escherichia coli*, *Pseudomonas aeruginosa*, *Enterococcus faecalis*, *Proteus vulgaris*, *Staphylococcus aureus*, *Acinetobacter baumannii*, CAMHB was supplemented with 2.5%~5.0% LHB for *Streptococcus*).

### 2.4 *Experiment on the evaluation of different combined effects*

The antibiotic sensitivity of bacterial strains was assessed according to Clinical Laboratory Standards Institute (CLSI) document M100-S24 (Wayne, 2014). MICs of antibiotics, flavonoids of *Ilex rotunda Thunb*. and their combinations were determined using the microdilution checkerboard method. *Escherichia coli*, *Pseudomonas aeruginosa*, *Enterococcus faecalis*, *Proteus vulgaris*, *Staphylococcus aureus*, *Acinetobacter baumannii* were tested in MH (pH=7.3), while CAMHB was supplemented with 2.5%~5.0% LHB for cultivation of *Streptococcus*.

Microdilution techniques were used to test the interactions between total flavonoids and other antibiotics. All these methods were developed for the detection of drug interactions, for there is no standardized method known to evaluate interaction between total flavonoids and antibiotics. Dilution procedures were performed according to CLSI protocol M7A7-2006(Adrar, Oukil & Bedjou, 2016).

Serial dilutions of total flavonoids and antibiotics were prepared, different combinations of these antibacterial were made and tested. Each well of the plate contain different concentrations of flavonoids and antibiotics which was shown in Table 1. The final volume in each well was equal to 100μL. The microtiter plates were incubated at 37°C for 16-18h.

**Table 1.**
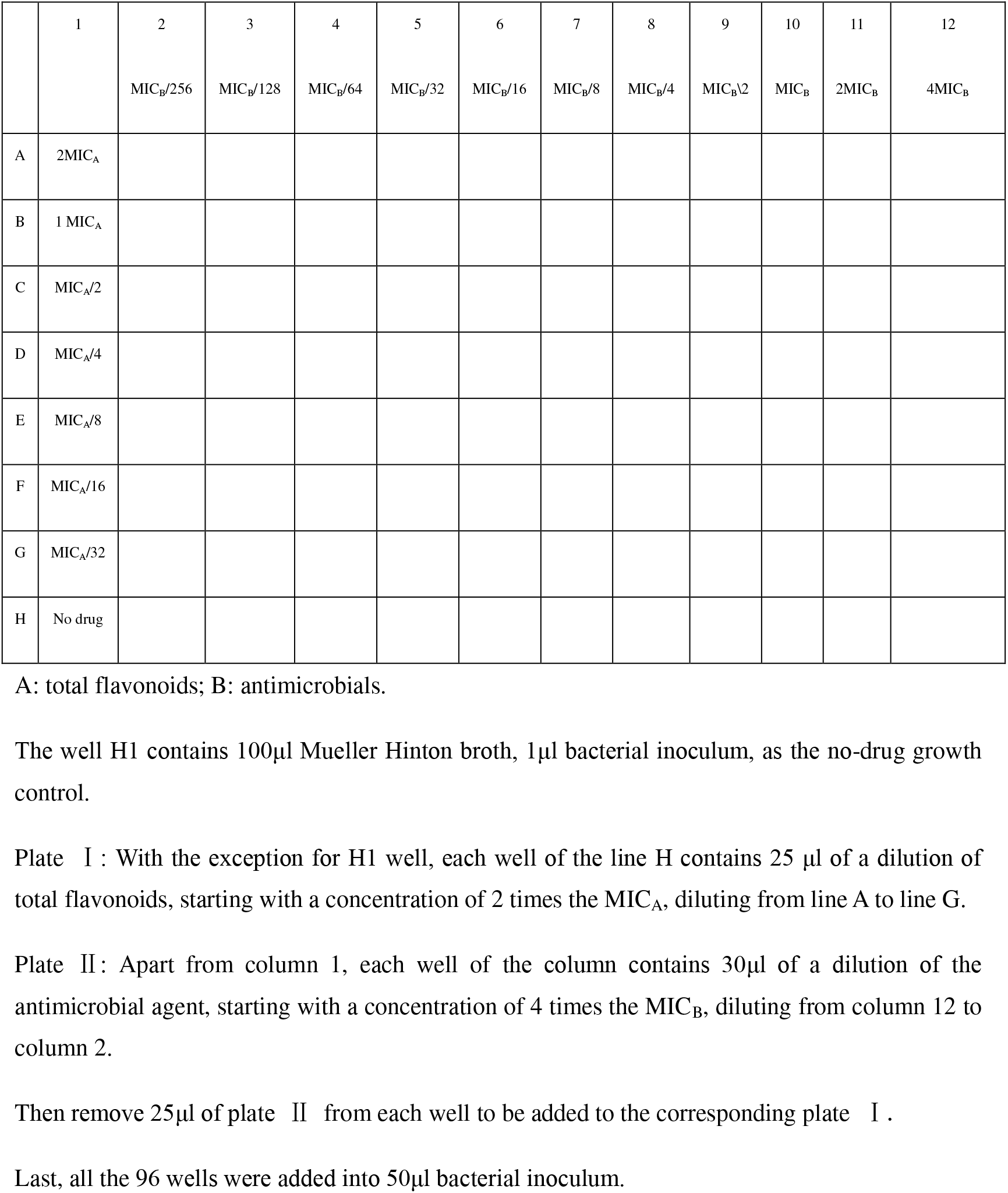
Microdilution technique used for the evaluation of combination effect on bacterial strain.

The FICI was calculated to evaluate the combined antimicrobial effect of antibiotics and flavonoids:

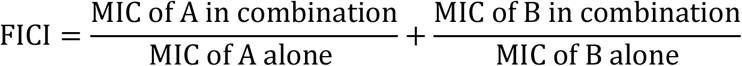

In the present study, the combined antibacterial effects of antibiotics and flavonoids were synergistic effect, addition, indifferent effect or antagonistic effect when the results were FICI ≤ 0.5, 0.5 < FICI ≤1, 1 < FICI ≤ 2 and FICI > 2, respectivel(Kurek, Nadkowska, Pliszka & Wolska, 2012).

## 3. Results

### 3.1 *Antibacterial activities of total flavonoids on different drug-resistant bacteria*

The MIC values of total flavonoids on various bacteria isolated including gram positive bacteria and gram negative bacteria were shown in Table 2. The minimum inhibitory concentration of total flavonoids against *Acinetobacter Streptococcus* and *Acinetobacter baumannii* were 20mg/ml, that of *Enterococcus faecalis*, *Proteus vulgaris*, *Staphylococcus aureus* were 40mg/ml. The MIC values of total flavonoids on *E.coli* and *Pseudomonas aeruginosa* were 80mg/ml, showing antibacterial effects.

**Table 2.** Minimum inhibitory concentrations (MICs; mg/mL) of total flavonoids from *Ilex Rotunda Thunb*.

| Bacteria | GRAM | MIC |
| --- | --- | --- |
| *Staphylococcus aureus* (R) | Positive | 40 |
| *Streptococcus* (R) | Positive | 20 |
| *Enterococcus faecalis* (R) | Positive | 40 |
| *Pseudomonas aeruginosa* (R) | Negative | 80 |
| *Proteus vulgaris* (R) | Negative | 40 |
| *Escherichia coli* (R) | Negative | 80 |
| *Acinetobacter Bauman* (R) | Negative | 20 |

R: resistant bacteria

### 3.2 *Antimicrobial activities of different combinations on gram negative bacteria*

Antimicrobial activities of different combinations on gram negative bacteria are shown in Table 3. Drug sensitivity test showed that *E.coli*, *Pseudomonas aeruginosa*, *Proteus vulgaris*, *Acinetobacter baumannii* were all multidrug-resistant bacteria. The effect of tested combinations was evaluated by MIC and FICI.

**Table 3.**
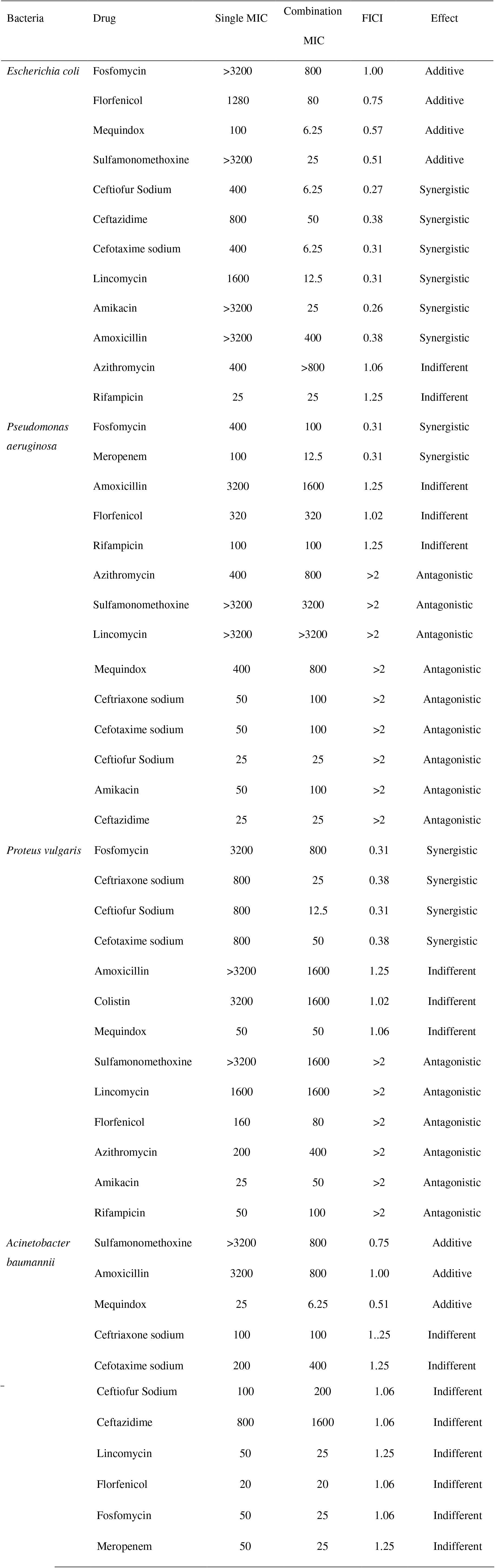
Minimum inhibitory concentrations (MICs; μg/mL) of antibiotics alone or with total flavonoids from *Ilex Rotunda Thunb*. against gram negative bacteria.

As shown in Table 3, an additive effect is obtained against *E.coli* by using flavonoids with fosfomycin, florfenicol, mequindox, sulfamonomethoxine, a synergistic effect is obtained against *E.coli* by using the combination flavonoids with ceftiofur sodium, ceftazidime, cefotaxime sodium, lincomycin, amikacin, amoxicillin, an indifferent effect is shown by using flavonoids/azithromycin and flavonoids/rifampicin combination against *Escherichia coli*.

For *Pseudomonas. aeruginosa*, the combination of flavonoids and fosfomycin, meropenem shows synergistic effect, an indifferent effect is shown by using flavonoids and amoxicillin, florfenicol, rifampicin combination, while using flavonoids with azithromycin sulfamonomethoxine, lincomycin, mequindox, ceftriaxone sodium, cefotaxime sodium shows antagonistic effects.

To *Proteus vulgaris*, the combination of flavonoids and fosfomycin, ceftriaxone sodium, ceftiofur Sodium, cefotaxime sodium shows synergistic effect, showing indifferent with amoxicillin, colistin, mequindox, an antagonistic effect is obtained with the following combination of flavonoids with sulfamonomethoxine, lincomycin, florfenicol, azithromycin, amikacin, rifampicin, respectively.

For *Acinetobacter baumannii*, an additive effect is obtained by using flavonoids with sulfamonomethoxine, amoxicillin, mequindox, showing indifference with ceftriaxone sodium, cefotaxime sodium, ceftiofur Sodium, ceftazidime, lincomycin, Florfenicol, fosfomycin, meropenem.

### 3.3 *Antimicrobial activities of different combinations on gram positive bacteria*

Antimicrobial activities of different combinations on gram positive bacteria are shown in Table 4. Drug sensitivity test showed that *Enterococcus faecalis*, *Streptococcus*, *Staphylococcus aureus* were all multidrug-resistant bacteria. The effect of tested combinations was evaluated by MIC and FICI.

As shown in Table 4, an additive effect is obtained against *Enterococcus faecalis* by using flavonoids with amoxicillin, florfenicol, rifampicin, a synergistic effect shown in the combination of flavonoids with amikacin, ceftazidime, ceftiofur Sodium, cefotaxime sodium, meropenem, ceftriaxone sodium, showing indifference by using flavonoids with sulfamonomethoxine, and an antagonistic effect is obtained by using flavonoids with colistin, lincomycin and azithromycin.

**Table 4.**
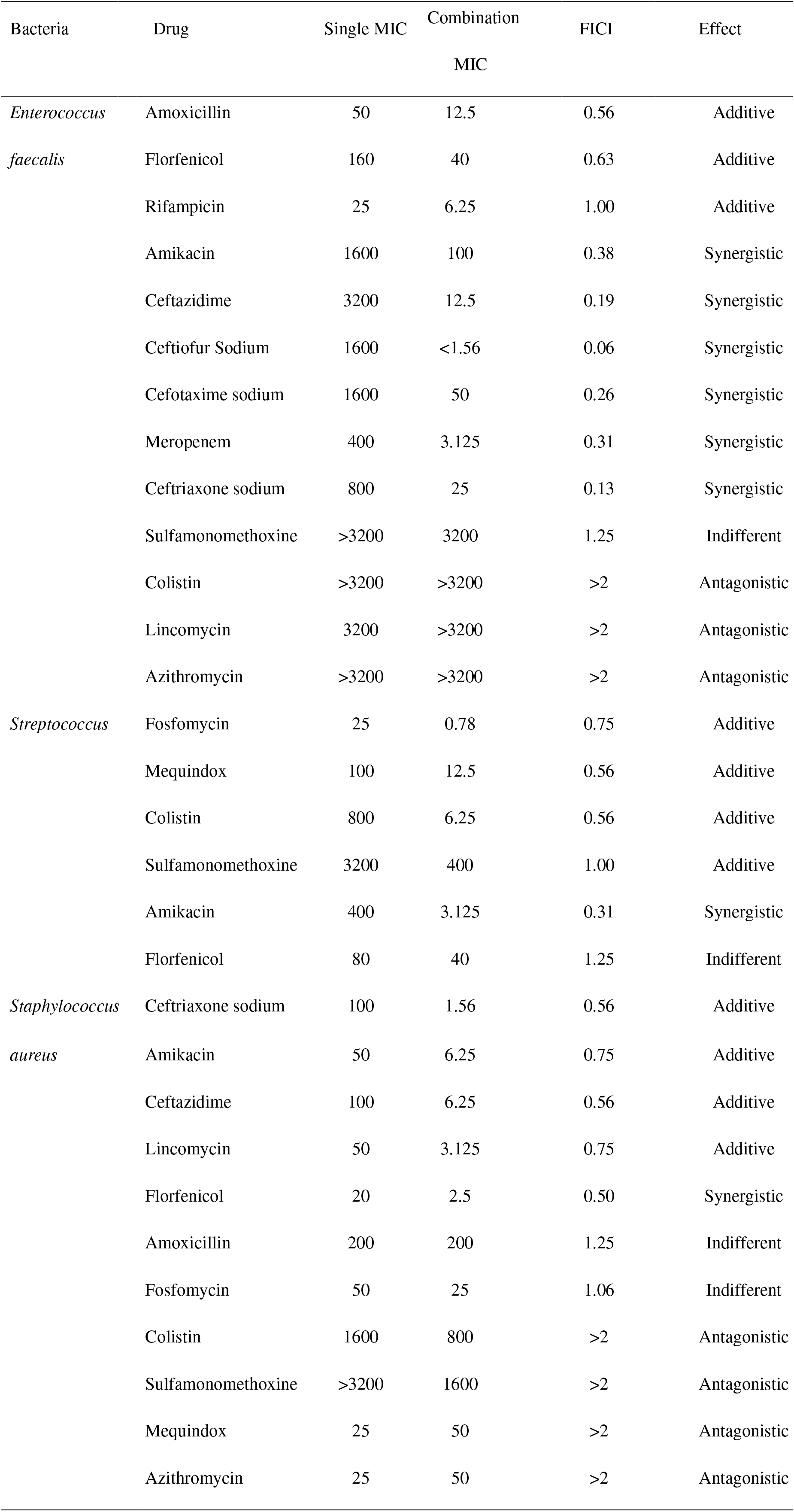
Minimum inhibitory concentrations (MICs; μg/mL) of antibiotics alone or with total flavonoids from *Ilex Rotunda Thunb*. against *gram positive bacteria*.

For *Streptococcus*, the combination of flavonoids with fosfomycin, mequindox, colistin, sulfamonomethoxine show additive action, showing synergistic effect by using flavonoids/amikacin combination, an indifferent effect is obtained with the combination of flavonoids/florfenicol.

An additive effect is obtained against *Staphylococcus aureus* by using the combination of flavonoids and ceftriaxone sodium, amikacin, ceftazidime, lincomycin, showing synergistic effect with florfenicol, the combination of flavonoids and amoxicillin, fosfomycin show indifference. In another hand, an antagonistic effect is obtained with the combination of flavonoids and colistin, sulfamonomethoxine, mequindox, azithromycin.

## 4. Discussion

The most relevant result of this study is that it demonstrates total flavonoids of *Ilex rotunda Thunb*. are equally effective against the above 7 different multidrug-resistant bacteria. Furthermore, the combinations of flavonoids and different antimicrobial agents are also effective against those multidrug-resistant bacteria.

Multidrug resistance has become a serious problem in the world, an urgent need for research into novel antibacterial agent and the development of efficacious combinations against multidrug-resistant bacterial clinical isolates are becoming more and more important(Oliveira et al., 2017; Bisi-Johnson, Obi, Samuel, Eloff & Okoh, 2017). Plant extracts used in combination with the antibiotic can be water extract, extracts of different solvents in different parts of plant or a serum containing plant extracts (Ahmad & Aqil, 2007). Compared with herbal medicine, antibiotics are easier to cause the problem of bacterial resistance due to their composition is clear and single while these herbal medicine own complex and varied composition (Schwarz, Kehrenberg & Walsh, 2001; Mishra, Rath, Swain, Ghosh, Das & Padhy, 2017). Medicinal plants have antibacterial action in vitro, enhance the antibiotic activity(Barreto et al., 2016), delay or eliminate bacterial tolerance(Roy-Leon, Lauzon, Toye, Singhal & Cameron, 2005), while because they contain a variety of effective ingredients which are not easy to be separated completely, so these medicinal plants cannot easily replace antibiotics. But the combined use of them and antibiotics may be the development trend of the clinical application of Chinese medicine.

In general, the target of antibiotics is single, but bacterial mutation is happening more and more, it is difficult to avoid the resistance mutations in bacteria(Schwarz, Kehrenberg & Walsh, 2001). One of TCM antibacterial characteristics is antibacterial ingredients are varied(Xiong, Li, Wang, Hong, Tang & Luo, 2013), which can inhibit or kill bacteria in different ways at the same time(Khan et al., 2015), so the choice of TCM combined with antibiotics can not only effect for a longer time, have a broad spectrum of antibacterial effect(Bisi-Johnson, Obi, Samuel, Eloff & Okoh, 2017d; Rajakumar, Gomathi, Thiruvengadam, Devi, Kalpana & Chung, 2017; Teanpaisan, Kawsud, Pahumunto & Puripattanavong, 2017), but also can be used as inhibitors of multidrug resistance associated protein targets, promoting antibiotics play a better role, reducing the dosage of antibiotics, further reducing the possibility of bacterial resistance to antibiotics, and providing certain clinical medication safety the security(Eumkeb, Tanphonkrang, Sirichaiwetchakoon, Hengpratom & Naknarong, 2017; Eumkeb, Siriwong & Thumanu, 2012; Eumkeb, Siriwong, Phitaktim, Rojtinnakorn & Sakdarat, 2012).

The experiment results proved that the total flavonoids of *Ilex rotunda Thunb*. combined with different antibacterial agents achieve a good effect, and screen combinations that can play a synergistic or additive inhibitory effect against different drug-resistant bacteria, providing a certain theoretical basis for the application and research and development of *Ilex rotunda Thunb*.

## 4 Conclusion

In this study, the results demonstrate that different combinations of flavonoids and *Ilex rotunda Thunb*. are equally effective against 7 multidrug-resistant bacteria isolated. Analyses show that flavonoids in combination with other antimicrobials exhibit different effects (additive, synergistic, indifferent and antagonistic) against drug-resistant bacteria. This is probably due to different mechanisms of action involved in this case. Flavonoids extracted in *Ilex rotunda Thunb*. showed high antibacterial activities. The study of these combinations show additive or synergistic action against drug-resistant bacteria could be useful to predict the efficacy of drugs in clinical development phases.

## 5 Conflict of interest statement

None of the authors of this paper have a financial or personal relationship with other people or organizations that could inappropriately influence or bias the content of the paper.

## 6 Acknowledgements

The authors gratefully acknowledge the support by the National Science Foundation of China (31460675).

## References

Adrar, N., Oukil, N., & Bedjou, F. (2016). Antioxidant and antibacterial activities of Thymus numidicus and Salvia officinalis essential oils alone or in combination. Industrial Crops and Products, 88, 112–119.

Ahmad, I., & Aqil, F. (2007). In vitro efficacy of bioactive extracts of 15 medicinal plants against ESbetaL-producing multidrug-resistant enteric bacteria. Microbiological Research, 162(3), 264–275.

Al-Alawi, R. A., Al-Mashiqri, J. H., Al-Nadabi, J., Al-Shihi, B. I., & Baqi, Y. (2017). Date Palm Tree (Phoenix dactylifera L.): Natural Products and Therapeutic Options. Frontiers in Plant Science, 8, 845.

Barbieri, R., Coppo, E., Marchese, A., Daglia, M., Sobarzo-Sanchez, E., Nabavi, S. F., & Nabavi, S. M. (2017). Phytochemicals for human disease: An update on plant-derived compounds antibacterial activity. Microbiological Research, 196, 44–68.

Barreto, H. M., Coelho, K. M., Ferreira, J. H., Dos, S. B., de Abreu, A. P., Coutinho, H. D., Da, S. R., de Sousa, T. O., Cito, A. M., & Lopes, J. A. (2016). Enhancement of the antibiotic activity of aminoglycosides by extracts from Anadenanthera colubrine (Vell.) Brenan var. cebil against multi-drug resistant bacteria. Natural Product Research, 30(11), 1289–1292.

Bisi-Johnson, M. A., Obi, C. L., Samuel, B. B., Eloff, J. N., & Okoh, A. I. (2017a). Antibacterial activity of crude extracts of some South African medicinal plants against multidrug resistant etiological agents of diarrhoea. BMC Complement Altern Med, 17(1), 321.

Bisi-Johnson, M. A., Obi, C. L., Samuel, B. B., Eloff, J. N., & Okoh, A. I. (2017d). Antibacterial activity of crude extracts of some South African medicinal plants against multidrug resistant etiological agents of diarrhoea. BMC Complement Altern Med, 17(1), 321.

Eskandary, S., Tahmourespour, A., Hoodaji, M., & Abdollahi, A. (2017). The synergistic use of plant and isolated bacteria to clean up polycyclic aromatic hydrocarbons from contaminated soil. J Environ Health Sci Eng, 15, 12.

Eumkeb, G., Siriwong, S., Phitaktim, S., Rojtinnakorn, N., & Sakdarat, S. (2012). Synergistic activity and mode of action of flavonoids isolated from smaller galangal and amoxicillin combinations against amoxicillin-resistant Escherichia coli. Journal of Applied Microbiology, 112(1), 55–64.

Eumkeb, G., Siriwong, S., & Thumanu, K. (2012). Synergistic activity of luteolin and amoxicillin combination against amoxicillin-resistant Escherichia coli and mode of action. J Photochem Photobiol B, 117, 247–253.

Eumkeb, G., Tanphonkrang, S., Sirichaiwetchakoon, K., Hengpratom, T., & Naknarong, W. (2017). The synergy effect of daidzein and genistein isolated from Butea superba Roxb. on the reproductive system of male mice. Natural Product Research, 31(6), 672–675.

Jones, M. E., Draghi, D. C., Thornsberry, C., Karlowsky, J. A., Sahm, D. F., & Wenzel, R. P. (2004). Emerging resistance among bacterial pathogens in the intensive care unit--a European and North American Surveillance study (2000-2002). Ann Clin Microbiol Antimicrob, 3, 14.

Khan, I., Ahmad, K., Khalil, A. T., Khan, J., Khan, Y. A., Saqib, M. S., Umar, M. N., & Ahmad, H. (2015). Evaluation of antileishmanial, antibacterial and brine shrimp cytotoxic potential of crude methanolic extract of Herb Ocimum basilicum (Lamiacea). Journal of Traditional Chinese Medicine, 35(3), 316–322.

Kurek, A., Nadkowska, P., Pliszka, S., & Wolska, K. I. (2012). Modulation of antibiotic resistance in bacterial pathogens by oleanolic acid and ursolic acid. Phytomedicine, 19(6), 515–519.

Liu, W., Peng, Y., Chen, H., Liu, X., Liang, J., & Sun, J. (2017). Triterpenoid Saponins with Potential Cytotoxic Activities from the Root Bark ofIlex rotundaThunb. Chemistry & Biodiversity, 14(2), e1600209.

Liu, Y., Cheng, Y., Yang, H., Hu, L., Cheng, J., Ye, Y., & Li, J. (2017). Characterization of Extended-Spectrum beta-Lactamase Genes of Shigella flexneri Isolates With Fosfomycin Resistance From Patients in China. Annals of Laboratory Medicine, 37(5), 415–419.

Miravitlles, M., & Anzueto, A. (2017). Chronic Respiratory Infection in Patients with Chronic Obstructive Pulmonary Disease: What Is the Role of Antibiotics? International Journal of Molecular Sciences, 18(7).

Mishra, M. P., Rath, S., Swain, S. S., Ghosh, G., Das, D., & Padhy, R. N. (2017). In vitro antibacterial activity of crude extracts of 9 selected medicinal plants against UTI causing MDR bacteria. Journal of King Saud University-Science, 29(1), 84–95.

Naz, R., Ayub, H., Nawaz, S., Islam, Z. U., Yasmin, T., Bano, A., Wakeel, A., Zia, S., & Roberts, T. H. (2017a). Antimicrobial activity, toxicity and anti-inflammatory potential of methanolic extracts of four ethnomedicinal plant species from Punjab, Pakistan. BMC Complement Altern Med, 17(1), 302.

Oliveira, F. S., Freitas, T. S., Cruz, R., Costa, M., Pereira, R., Quintans-Junior, L. J., Andrade, T. A., Menezes, P., Sousa, B., Nunes, P. S., Serafini, M. R., Menezes, I., Araujo, A., & Coutinho, H. (2017). Evaluation of the antibacterial and modulatory potential of alpha-bisabolol, beta-cyclodextrin and alpha-bisabolol/beta-cyclodextrin complex. Biomedicine & Pharmacotherapy, 92, 1111–1118.

Rajakumar, G., Gomathi, T., Thiruvengadam, M., Devi, R. V., Kalpana, V. N., & Chung, I. M. (2017). Evaluation of anti-cholinesterase, antibacterial and cytotoxic activities of green synthesized silver nanoparticles using from Millettia pinnata flower extract. Microb Pathog, 103, 123–128.

Ramezani, M., Rahmani, F., & Dehestani, A. (2017). Comparison between the effects of potassium phosphite and chitosan on changes in the concentration of Cucurbitacin E and on antibacterial property of Cucumis sativus. BMC Complement Altern Med, 17(1), 295.

Roy-Leon, J. E., Lauzon, W. D., Toye, B., Singhal, N., & Cameron, D. W. (2005). In vitro and in vivo activity of combination antimicrobial agents on Haemophilus ducreyi. J Antimicrob Chemother, 56(3), 552–558.

Santiago, C., Pang, E. L., Lim, K. H., Loh, H. S., & Ting, K. N. (2015). Inhibition of penicillin-binding protein 2a (PBP2a) in methicillin resistant Staphylococcus aureus (MRSA) by combination of ampicillin and a bioactive fraction from Duabanga grandiflora. BMC Complement Altern Med, 15, 178.

Schwarz, S., Kehrenberg, C., & Walsh, T. R. (2001). Use of antimicrobial agents in veterinary medicine and food animal production. Int J Antimicrob Agents, 17(6), 431–437.

Soudeiha, M., Dahdouh, E. A., Azar, E., Sarkis, D. K., & Daoud, Z. (2017). In vitro Evaluation of the Colistin-Carbapenem Combination in Clinical Isolates of A. baumannii Using the Checkerboard, Etest, and Time-Kill Curve Techniques. Front Cell Infect Microbiol, 7, 209.

Teanpaisan, R., Kawsud, P., Pahumunto, N., & Puripattanavong, J. (2017). Screening for antibacterial and antibiofilm activity in Thai medicinal plant extracts against oral microorganisms. J Tradit Complement Med, 7(2), 172–177.

Usman, A. M., Khurram, M., Khan, T. A., Faidah, H. S., Ullah, S. Z., Ur, R. S., Haseeb, A., Ilyas, M., Ullah, N., Umar, K. S., & Iriti, M. (2016). Effects of Luteolin and Quercetin in Combination with Some Conventional Antibiotics against Methicillin-Resistant Staphylococcus aureus. International Journal of Molecular Sciences, 17(11).

Wayne, P. (2014). CLSI. Performance Standards for Antimicrobial Susceptibility Testing;Twenty-Fourth Informational Supplement., CLSI document M100-S24: Clinical and Laboratory Standards Institute.

Xiong, J., Li, S., Wang, W., Hong, Y., Tang, K., & Luo, Q. (2013). Screening and identification of the antibacterial bioactive compounds from Lonicera japonica Thunb. leaves. Food Chemistry, 138(1), 327–333.

Ye, J., Cheng, H., Li, H., Yang, Y., Zhang, S., Rauf, A., Zhao, Q., & Ning, G. (2017). Highly synergistic antimicrobial activity of spherical and flower-like hierarchical titanium dioxide/silver composites. J Colloid Interface Sci, 504, 448–456.

Zakaryan, H., Arabyan, E., Oo, A., & Zandi, K. (2017). Flavonoids: promising natural compounds against viral infections. Archives of Virology.

Zou, L., Lu, J., Wang, J., Ren, X., Zhang, L., Gao, Y., Rottenberg, M. E., & Holmgren, A. (2017). Synergistic antibacterial effect of silver and ebselen against multidrug-resistant Gram-negative bacterial infections. EMBO Molecular Medicine.

